# Association Mapping from Sequencing Reads Using *K*-mers

**DOI:** 10.1101/141267

**Authors:** Atif Rahman, Ingileif Hallgrímsdóttir, Michael B. Eisen, Lior Pachter

**Affiliations:** Department of Electrical Engineering and Computer Sciences, University of California, Berkeley, California, United States of America; Department of Statistics, University of California, Berkeley, California, United States of America; Department of Molecular & Cell Biology, University of California, Berkeley, California, United States of America; Howard Hughes Medical Institute, University of California, Berkeley, California, United States of America; Department of Mathematics, University of California, Berkeley, California, United States of America

## Abstract

Genome wide association studies (GWAS) rely on microarrays, or more recently mapping of whole-genome sequencing reads, to genotype individuals. The reliance on prior sequencing of a reference genome for the organism on which the association study is to be performed limits the scope of association studies, and also precludes the identification of differences between cases and controls outside of the reference. We present an alignment free method for association studies that is based on counting k-mers in sequencing reads, testing for associations directly between k-mers and the trait of interest, and local assembly of the statistically significant k-mers to identify sequence differences. Results with simulated data and an analysis of the 1000 genomes data provide a proof of principle for the approach. In a pairwise comparison of the Toscani in Italia (TSI) and the Yoruba in Ibadan, Nigeria (YRI) populations we find that sequences identified by our method largely agree with results obtained using standard GWAS based on variant calling from mapped reads. However unlike standard GWAS, we find that our method identifies associations with structural variations and sites not present in the reference genome revealing sequences absent from the human reference genome. We also analyze data from the Bengali from Bangladesh (BEB) population to explore possible genetic basis of high rate of mortality due to cardiovascular diseases (CVD) among South Asians and find significant differences in frequencies of a number of non-synonymous variants in genes linked to CVDs between BEB and TSI samples, including the site rs1042034, which has been associated with higher risk of CVDs previously, and the nearby rs676210 in the *Apolipoprotein B (ApoB)* gene.

**Author Summary:** We present a method for associating regions in genomes to traits or diseases. The method is based on finding differences in frequencies of short strings of letters in sequencing reads and do not require reads to be aligned to a reference genome. This makes it applicable to study of organisms with no or incomplete reference genomes. We test our method with simulated data and sequencing data from the 1000 genomes project and find agreement with the conventional approach based on alignment to a reference genome. In addition, our method finds associations with sequences not in reference genomes and reveals sequences missing from the human reference genome. We also explore high rates of mortality due to cardiovascular diseases among South Asians and find prevalence of variations in genes associated with heart diseases in samples from the Bengali from Bangladesh population including one that has been reported to be associated with early onset of cardiovascular diseases.

## Introduction

Association mapping refers to the linking of genotypes to phenotypes. Most often this is done using a genome-wide association study (GWAS) with single nucleotide polymorphisms (SNPs). Individuals are genotyped at a set of known SNP locations using a SNP array. Then each SNP is tested for statistically significant association with the phenotype. In recent years thousands of genome-wide association studies have been performed and regions associated with traits and diseases have been located.

However, this approach has a number of limitations. First, designing SNP arrays requires knowledge about the genome of the organism and where the SNPs are located in the genome. This makes it hard to apply to study organisms other than human. Even the human reference genome is incomplete [1] and association mapping to regions not in the reference is difficult. Second, structural variations such as insertion-deletions (indels) and copy number variations are usually ignored in these studies. Despite the many GWA studies that have been performed a significant amount of heritability is yet to be explained. This is known as the “missing heritability” problem [2]. A hypothesis is some of the missing heritability is due to structural variations. Third, the phenotype might be caused by rare variants which are not on the SNP chip. In last two cases, follow up work is required to find the causal variant even if association is detected in the GWAS.

Some of these limitations can be overcome by utilizing high throughput sequencing data. As sequencing gets cheaper association mapping using next generation sequencing is becoming feasible. The current approach to doing this is to map all the reads to a reference genome followed by variant calling. Then these variants can be tested for association. But this again requires a reference genome and it may induce biases in variant calling and regions not in the reference genome will not be included in the study. Moreover, sequencing errors make genotype calling difficult when sequencing depth is low [3] and in repetitive regions. Methods have been proposed to do population genetics analyses that avoid the genotype calling step [4, 5] but these methods still require reads to be aligned to a reference genome. An alternate approach is simultaneous *de novo* assembly and genotyping using a tool such as Cortex [6] but this is not suited to large number of individuals. Furthermore, both these approaches are computationally very expensive.

In the past, alignment free methods have been developed for a number of problems including transcript abundance estimation [7], sequence comparison [8], phylogeny estimation [9], etc. Nordstrom *et al.* introduced a pipeline called needle in the *k*-stack (NIKS) for mutation identification by comparison of sequencing data from two strains using *k*-mers [10]. Here we present an alignment free method for association mapping. It is based on counting *k*-mers and identifying *k*-mers associated with the phenotype. The overlapping k-mers found are then assembled to obtain sequences corresponding to associated regions. Our method is applicable to association studies in organisms with no or incomplete reference genome. Even if a reference genome is available, this method has the advantage of avoiding aligning and genotype calling thus allowing association mapping to many types of varaints using the same pipeline and to regions not in the reference.

We have implemented our method in a software called ‘hitting associations with k-mers’ (HAWK). Experiments with simulated and real data demonstrate the promises of this approach. We leave taking into account confounding factors such as population structure as future work and apply our method to analyze sequencing data from three populations in the 1000 genomes project treating population identity as the trait of interest. Agreement with sites found using read alignment and genotype calling indicate that k-mer based association mapping will be applicable to studying disease associations.

## Methods

### Association mapping with *k*-mers

We present a method for finding regions associated with a trait using sequencing reads without mapping reads to reference genomes. The workflow is illustrated in Fig 1. Given sequencing reads from case and control samples, we count k-mers appearing in each sample. We assume the counts are Poisson distributed and test k-mers for statistically significant association with case or control using likelihood ratio test for nested models (see Supplementary for details). The differences in k-mer counts may be due to single nucleotide polymorphisms (SNPs), insertion-deletions (indels) and copy number variations. The k-mers are then assembled to obtain sequences corresponding to each region.

### Counting *k*-mers

The first step in our method for association mapping from sequencing reads using k-mers is to count k-mers in sequencing reads from all samples. To count k-mers we use the multi-threaded hash based tool Jellyfish developed by Marcais and Kingsford [11]. We use k-mers of length 31 and ignore k-mers that appear once in a sample for computational and memory efficiency as they are likely from sequencing errors.

### Finding significant *k*-mers

Then for each k-mer we test whether that k-mer appears significantly more times in case or control datasets compared to the other using a likelihood ratio test for nested models. Suppose, a particular *k*-mer appears *K*_1_ times in cases and *K*_2_ times in controls, and *N*_1_ and *N*_2_ are the total number of *k*-mers in cases and controls respectively. The *k*-mer counts are assumed to be Poisson distributed with rates *θ*_1_ and *θ*_2_ in cases and controls. The null hypothesis is *H*_0_: *θ*_1_ = *θ*_2_ = *θ* and the alternate hypothesis is *H*_1_: *θ*_1_ ≠ *θ*_2_. The likelihoods under the alternate and the null are given by (see Supplementary for details)

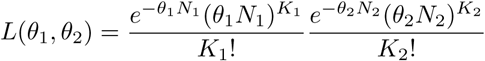

and

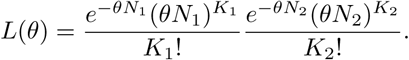

Since the null model is a special case of the alternate model, 2 ln Λ is approximately chi-squared distributed with one degree of freedom where Λ is the likelihood ratio. We get a p-value for each k-mer using the approximate *χ*^2^ distribution of the likelihood ratio and perform Bonferroni corrections to account for multiple testing.

**Figure 1.**
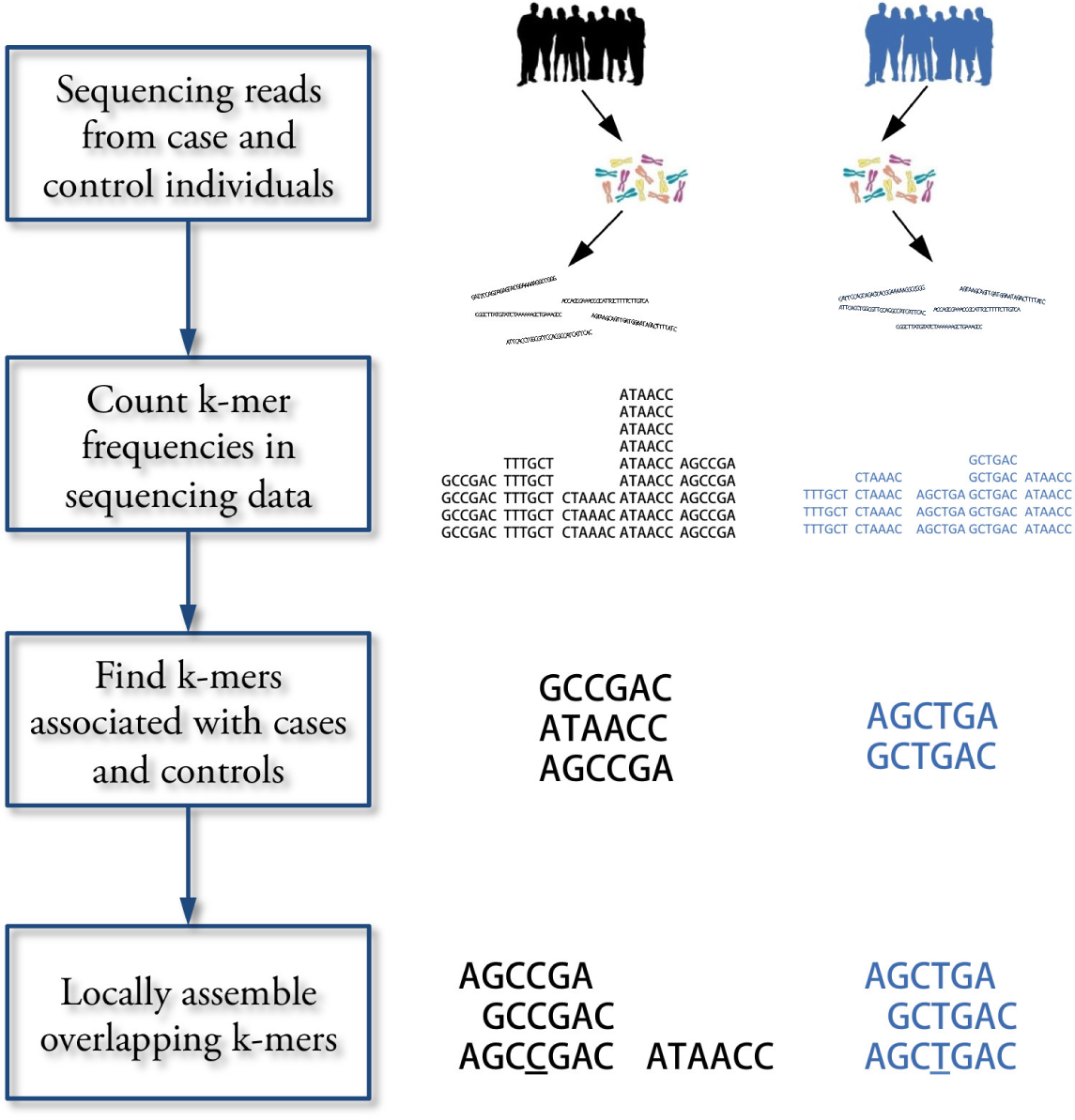
Workflow for association mapping using *k*-mers. The Hawk pipeline starts with sequencing reads from two sets of samples. The first step is to count k-mers in reads from each sample. Then k-mers with significantly different counts in two sets are detected using likelihood ratio test. Finally, overlapping k-mers are assembled into sequences to get one or few sequences for each associated locus.

### Merging *k*-mers

We then merge overlapping k-mers to get a sequence for each differential site using the assembler ABySS [12]. ABySS was used as the assemblies it generated were found to cover more of the sequences to be assembled compared to other assemblers [13]. We construct the de Bruijn graph using hash length of 25 and retain assembled sequences of length at least 49. It is also possible to merge k-mers and pair sequences from cases and controls using the NIKS pipeline [10]. However, we find that this is time consuming when we have many significant k-mers. Moreover, when number of cases and controls are not very high we do not have enough power to get both of sequences to be paired and as such pairing is not possible.

### Implementation

Our method is implemented in a tool called ‘hitting associations with k-mers’ (HAWK) using C++. To speed up the computation we use a multi-threaded implementation. In addition, it is not possible to load all the k-mers into memory at the same time for large genomes. So, we sort the k-mers and load them into memory in batches. To make the sorting faster Jellyfish has been modified to output internal representation of k-mers instead of the k-mer strings. In future the sorting step may be avoided by utilizing the internal ordering of Jellyfish or other tools for k-mer counting. The implementation is available at http://atifrahman.github.io/HAWK/

### Downstream analysis

The sequences can then be analyzed by aligning to a reference if one is available or by running BLAST [14] to check for hits to related organisms. The intersection results in this paper were obtained by mapping them to the human reference genome version GRCh37 using Bowtie2 [15] to be consistent with co-ordinates of genoptypes called by 1000 genomes project. The breakdown analysis was performed by first mapping to the latest version of the reference, hg38 and then running BLAST on some of the ones that did not map. Specific loci of interest were checked by aligning them to RefSeq mRNAs using Bowtie 2 and on the UCSC Human Genome Browser [16] by running BLAT [17].

## Results

### Verification with simulated data

The implementation was tested by simulating reads from genome of an *Escherichia coli* strain. We introduced different types mutations - single nucleotide changes, short indels (less than 10bp) and long indels (between 100bp and 1000bp) into the genome. Then `wgsim` of `SAMtools` [18] was used to first generate two sets of genomes by introducing more random mutations (both substitutions and indels) into the original and the modified genomes and then simulate reads with sequencing errors. The Hawk pipeline was then run on these two sets of sequencing reads. The fraction of mutations covered by resulting sequences are shown in S1 Fig) for varying numbers of case and control samples and different types of mutations. The results are consistent with calculation of power to detect k-mers for varying total k-mer coverage (S2 Fig) with slightly lower values expected due to sequencing errors and conditions imposed during assembly.

### Verification with 1000 genomes data

To analyze the performance of the method on real data we used sequencing reads from the 1000 genomes project [19]. The population identities were used as the phenotype of interest circumventing the need for correction of population structure. For verification, we used sequencing reads from 87 YRI individuals and 98 TSI individuals for which both sequencing reads and genotype calls were available at the time analysis was performed.

The analysis using k-mers resulted in 2,970,929 sequences associated with YRI samples and 1,865,285 sequences of significant association with TSI samples. We also performed similar analysis with genotype calls. VCFtools [20] was used to obtain number of individuals with 0, 1 and 2 copies of one of the alleles for each SNP site. Each site was then tested to check whether the allele frequencies are significantly different in two samples using likelihood ratio test for nested models for multinomial distribution (details in S1 Text). We found that 2,658,964 out of the 39,706,715 sites had allele frequencies that are significantly different.

Figure 2(a) shows the extent of overlap among these discarding the sequences that did not map to the reference. We find that 80.3% (2,135,415 out of 2,658,964) of the significant sites overlapped with some sequence found using Hawk. Approximately 95.2% of the sites overlapped with at least one k-mer.

We also observe that around 42% of sequences found using *k*-mers do not overlap with any sites found significant using genotype calling. While upto 20% of them correspond to regions for which we did not have genotype calls (chromosome Y, mitochondrial DNA and small contigs), repetitive regions where genotype calling is difficult and structural variations, many of the remaining sequences are possibly due to more power of the test based on counts than the one using only number of copies of an allele. We performed Monte Carlo simulations to determine powers of the two tests. Figure 2(b) shows the fraction of trials that passed the p-value threshold after Bonferroni correction as the allele frequencies in cases were increased keeping the allele frequencies of control fixed at 0.

**Figure 2.**
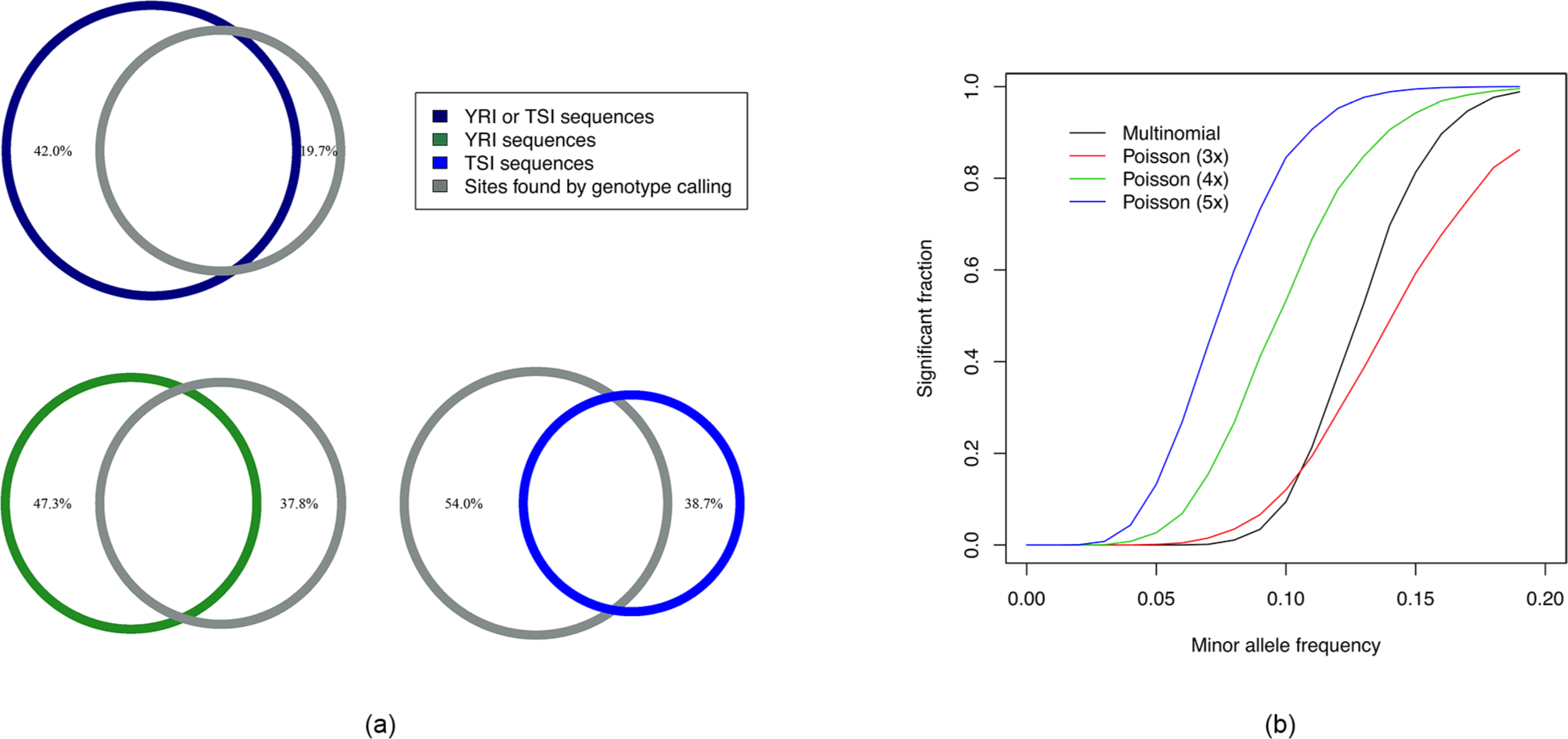
Intersection analysis and comparison of powers of tests. (a) Venn diagrams showing intersections among sequences obtained using Hawk and significant sites found by genotype calling. 80.3% of the sites overlapped with some sequence. Around 42% of sequences do not overlap with any such site which can be explained by more types of variants found by Hawk as well as more power of the test using Poisson compared to Multinomial distribution. (b) Fraction of runs found significant (after Bonferroni correction) by tests against minor allele frequency of the case samples (with that of the controls fixed at 0) are shown. The curves labeled multinomial and Poisson correspond to likelihood ratio test using multinomial distribution and Poisson distributions with different k-mer coverage.

This is consistent with greater fraction of sequences in YRI (47.3%) not overlapping with sites obtained by genotyping compared to TSI (38.7%) as some low frequency variations in African populations were lost in other populations due to population bottleneck during the migration out of Africa. However, some false positives may result due to discrepancies in sequencing depth of the samples and sequencing biases. We provide scripts to lookup number of individuals with constituent k-mers and leave dealing with these confounding factors as well as population structure as future work.

Table 1 shows p-values of some of the well known sites of variation between African and European populations.

### HAWK maps associations to different types of variants

Hawk enables mapping associations to different types of variants using the same pipeline. Figure 3(a) shows breakdown of types of variants found associated with YRI and TSI populations. The ‘Multiple SNPs/Structural’ entries correspond to sequences of length greater than 61 (the maximum length of a sequence due to a single SNP with k-mer size of 31). In addition to SNPs we find associations to sites with indels and structural variations. Furthermore, we find sequences that map to multiple regions in the genome indicating copy number of variations or sequence variation in repeated regions where genotype calling is known to be difficult. Although the majority of the sequences map outside of genes, we find variants in genes including in coding regions (Figure 3(b)).

**Table 1.** Known variants in YRI-TSI comparison.

| Gene | SNP ID | Description | Allele | p-value | %YRI | %TSI |
| --- | --- | --- | --- | --- | --- | --- |
| *ACKR1* | rs2814778 | Duffy antigen | C | $9.72 \times 10^{-114}$ | 84.39% | 1.78% |
| *SLC24A5* | rs1426654 | Skin pigmentation | G | $8.45 \times 10^{-144}$ | 87.39% | 1.02% |
| *SLC45A2* | rs16891982 | Skin/hair color | C | $1.89 \times 10^{-122}$ | 92.18% | 4.67% |
| *G6PD* | rs1050829 | G6PD deficiency | C | $1.53 \times 10^{-29}$ | 24.92% | 1.02% |
| *G6PD* | rs1050828 | G6PD deficiency | T | $5.83 \times 10^{-25}$ | 18.32% | 0.00% |

Table 1 shows p-values of sequences at some well known sites of variation between populations. The (%) values denote fraction of individuals in the sample with the allele present. The p-values and % values are averaged over $k$ -mers constituting the associated sequences.

**Figure 3.**
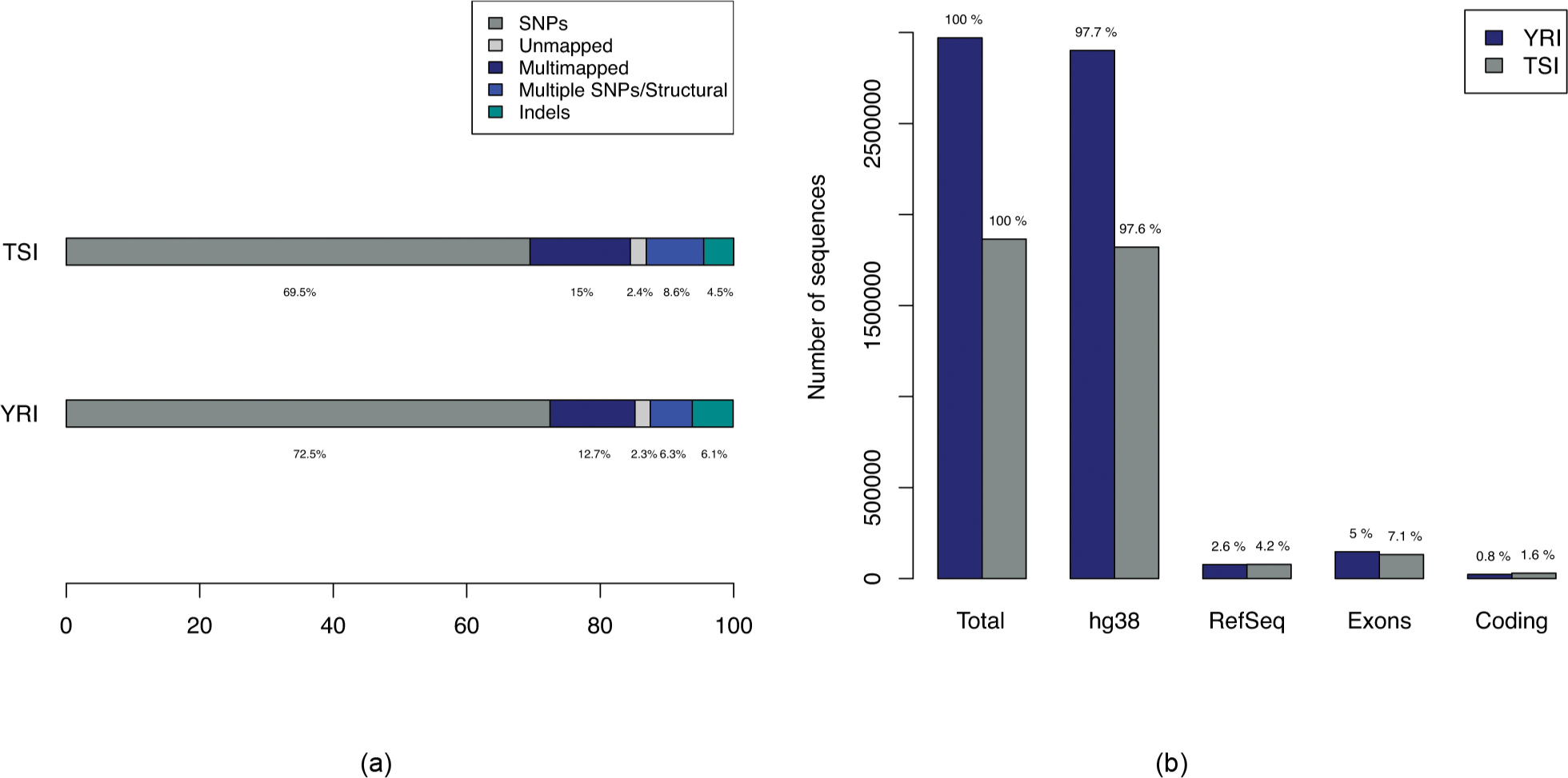
Breakdown of types of variations in comparison of YRI-TSI. (a) Bars showing breakdown of 2,970,929 and 1,865,285 sequences associated with YRI and TSI samples respectively. The ‘Multiple SNPs/Structural’ entries correspond to sequences of length greater than 61, the maximum length of a sequence due to a single SNP with k-mer size of 31 and ‘SNPs’ correspond to sequences of maximum length of 61. (b) Numbers of sequences with alignments to hg38, RefSeq mRNAs and Ensembl exons and coding regions.

We performed similar analysis on sequencing reads available from 87 BEB and 110 TSI individuals from the 1000 genomes project and obtained 529,287 and 462,122 sequences associated with BEB and TSI samples respectively, much fewer than the YRI-TSI comparison. S3 Fig shows breakdown of probable variant types corresponding to the sequences found associated with BEB and TSI samples.

Histograms of sequence lengths show (S4 Fig, S5 Fig) peaks at 61bp which is the maximum length corresponding to a single SNP for k-mer size of 31. We also see drops off after 98bp in all cases providing evidence for multinucleotide mutations (MNMs) reported in [21] since this is the maximum sequence length we can get when k-mers of size 31 are assembled with minimum overlap of 24.

### HAWK reveals sequences not in the human reference genome

As Hawk is an alignment free method for mapping associations, it is able to find associations in regions that are not in the human reference genome. The analysis resulted in 94,795 and 66,051 sequences of lengths up to 2,666bp and 12,467bp associated with YRI and TSI samples respectively that did not map to the human reference genome. Similarly BEB-TSI comparison yielded 19,584 and 18,508 sequences with maximum lengths of 1761bp and 2149bp associated with BEB and TSI respectively.

We found that few of the sequences associated with TSI samples some as long as 12kbp and 2kbp in comparisons against YRI and BEB respectively that mapped to the Epstein–Barr virus (EBV) genome, strain B95-8 [GenBank: V01555.2]. EBV strain B95-8 was used to transform B cells into lymphoblastoid cell lines (LCLs) in the 1000 Genomes Project and is a known contaminant in the data [22].

Table 2 summarizes the sequences that could not be mapped to either the human reference genome or the Epstein-Barr virus genome using Bowtie2. Although an exhaustive analysis of all remaining sequences using BLAST is difficult, we find sequences associated with YRI that do not map to the human reference genome (hg38) with high score but upon running BLAST aligned to other sequences from human (for example to [GenBank: AC205876.2] and some other sequences reported by Kidd *et al.* [23]). We also find sequences with no significant BLAST hits to human genomic sequences some with hits to closely related species. Similarly, we find sequences associated with TSI aligning to human sequences such as [GenBank: AC217954.1] not in the reference. Although there are much fewer long sequences obtained in the BEB-TSI comparison, we find sequences longer than 1kbp associated with each population with no BLAST hit.

**Table 2.** Summary of sequences not in the human reference genome.

| Population | Population compared to | Total no. sequences | No. sequences with length≥1000bp | Total length in sequences with length≥1000bp | No. sequences with length≥200bp | Total length in sequences with length≥200bp |
| --- | --- | --- | --- | --- | --- | --- |
| YRI | TSI | 94,795 | 41 | 59,956 | 478 | 225,426 |
| TSI | YRI | 66,051 | 10 | 13,896 | 184 | 77,383 |
| BEB | TSI | 19,584 | 3 | 3,835 | 75 | 33,954 |
| TSI | BEB | 18,508 | 2 | 2,105 | 81 | 28,134 |

Table 2 shows summary of sequences associated with different populations that did not map to the human reference genome (hg38) or to the Epstein-Barr virus genome.

### Differential prevalence of variants in genes linked to CVDs in BEB-TSI comparison

We noted that cardiovascular diseases (CVD) are a leading cause of mortality in Bangladesh [24] and age standardized death rates from CVDs in Bangladesh is higher compared to Italy [24]. Moreover, South Asians have high rates of acute myocardial infarction (MI) or heart failure at younger ages compared to other populations and in several countries migrants from South Asia have higher death rates from coronary heart disease (CHD) at younger ages compared with the local population [25, 26] and according to the Interheart Study, the mean age of MI among the poeple from Bangladesh is considerably lower than non-South Asians and the lowest among South Asians [27, 28]. This motivated us to explore probable underlying genetic causes.

The sequences of significant association with the BEB sample were aligned to RefSeq mRNAs and the ones mapping to genes linked to CVDs [29] were analyzed. Table 3 shows non-synonymous variants in such genes that are significantly more common in the BEB sample compared to the TSI sample. It is worth mentioning that the ‘C’ allele at the SNP site, rs1042034 in the gene *Apolipoprotein B (ApoB)* has been associated with increased levels of HDL cholesterol and decreased levels of Triglycerides [30] in individuals of European descent but individuals with the ‘CC’ genotype have been reported to have higher risk of CVDs in an analysis of the data from the Framingham Heart Study [31]. The SNP rs676210 has also been associated with a number of traits [32, 33]. Both alleles of higher prevalence in BEB at those sites have been found to be common in familial hypercholesterolemia patients in Taiwan [34]. On the other hand, prevalence of the risk allele, ‘T’ at rs3184504 in the gene *SH2B3* is higher in TSI samples compared to BEB samples.

We also observe a number of sites in the gene *Titin (TTN)* of differential allele frequencies in BEB and TSI samples (S1 Table). However, *TTN* codes for the largest known protein and although truncating mutations in *TTN* are known to cause dilated cardiomyopathy [35–37], no such effect of other kinds of mutations are known.

**Table 3.** Variants in genes linked to cardiovascular diseases.

| Gene | SNP ID | Variant type | Allele | p-value | %BEB | %TSI |
| --- | --- | --- | --- | --- | --- | --- |
| *APOB* | rs2302515 | Missense | C | $1.30 \times 10^{-12}$ | 29.29% | 8.37% |
| *APOB* | rs676210 | Missense | A | $7.73 \times 10^{-25}$ | 72.93% | 33.08% |
| *APOB* | rs1042034 | Missense | C | $2.28 \times 10^{-23}$ | 68.67% | 31.91% |
| *CYP11B2* | rs4545 | Missense | T | $1.31 \times 10^{-28}$ | 31.33% | 0.91% |
| *CYP11B1* | rs4534 | Missense | T | $9.36 \times 10^{-36}$ | 33.00% | 0.91% |
| *WNK4* | rs2290041 | Missense | T | $1.53 \times 10^{-14}$ | 13.24% | 0.47% |
| *WNK4* | rs55781437 | Missense | T | $1.30 \times 10^{-12}$ | 15.21% | 0.91% |
| *SLC12A3* | rs2289113 | Missense | T | $7.40 \times 10^{-13}$ | 8.14% | 0.00% |
| *SCNN1A* | rs10849447 | Missense | C | $8.67 \times 10^{-12}$ | 62.88% | 39.92% |
| *ABO* | - | 4bp (CTGT) deletion | - | $1.17 \times 10^{-13}$ | 29.15% | 10.55% |
| *ABO* | rs8176741 | Missense | A | $2.06 \times 10^{-16}$ | 27.70% | 8.45% |
| *SH2B3* | rs3184504 | Missense | C | $8.22 \times 10^{-23}$ | 92.88% | 63.87% |
| *RAI1* | rs3803763 | Missense | C | $1.32 \times 10^{-12}$ | 75.86% | 51.17% |
| *RAI1* | rs11649804 | Missense | A | $1.95 \times 10^{-19}$ | 81.57% | 52.79% |

Variants in genes linked to cardiovascular diseases found to be significantly more common in BEB sample compared to TSI sample. The (%) values denote fraction of individuals in the sample with the allele present. The p-values and % values are averaged over *k*-mers constituting the associated sequences.

## Discussion

In this paper, we presented an alignment free method for association mapping from sequencing reads. It is based on finding *k*-mers that appear significantly more times in one set of samples compared to the other and then locally assembling those *k*-mers. Since this method does not require a reference genome, it is applicable to association studies of organisms with no or incomplete reference genome. Even for human our method is advantageous as it can map associations in regions not in the reference or where variant calling is difficult.

We tested our method by applying it to data from the 1000 genomes project and comparing the results with the results obtained using the genotypes called by the project as well as using simulated data. We observe that more than 80% of the sites found using genotype calls are covered by some sequence obtained by our method while also mapping associations to regions not in the reference and in repetitive areas. Moreover, simulations suggest tests based on *k*-mer counts have more power than those based number of copies of an allele present at some site.

Breakdown analysis of the sequences found in pairwise comparison of YRI, TSI and BEB, TSI samples reveals that this approach allows mapping associations to SNPs, indels, structural and copy number variations through the same pipeline. In addition we find 2-4% of associated sequences are not present in the human reference genome some of which are longer than 1kbp. The YRI, TSI comparison yields almost 60kbp sequence associated with the YRI samples in sequences of length greater than 1kbp alone. This indicates populations around the world have regions in the genome not present in the reference emphasizing the importance of a reference free approach.

We explored variants in genes linked to cardiovascular diseases in the BEB, TSI comparison as South Asians are known to have a higher rate of mortality from heart diseases compared to many other populations. We find a number of non-synonymous mutations in those genes are more common in the BEB samples in comparison to the TSI ones underscoring the importance of association studies in diverse populations. The SNP rs1042034 in the gene *Apolipoprotein B (ApoB)* merits particular mention as the CC genotype at that site has been associated with higher risk of CVDs.

The results on simulated data and real data from the 1000 genomes project provide a proof of principle of this approach and motivate extension of this method to quantitative phenotypes and correction for population structure and other confounding factors and then application to association studies of disease phenotypes in humans and other organisms.

## Supporting Information

### S1 Text

#### Additional details

Contains additional details about the method and all supporting figures and tables.

### S1 Fig

#### Sensitivity with simulated *E. coli* data

The figure shows sensitivity for varying number of case and control samples for different types of mutations. Sensitivity is defined as the percentage of differing nucleotides that are covered by a sequence. All of the sequences covered some location of mutation.

### S2 Fig

#### Power for different k-mer coverages

The figure shows power to detect a k-mer present in all case samples and no control sample against total k-mer coverage of cases using Bonferroni correction for different number of total tests for p-value=0.05.

### S3 Fig

#### Breakdown of types of variations in BEB-TSI comparison

(a) Bar plots showing breakdown of 529,287 and 462,122 sequences associated with BEB and TSI samples respectively. The ‘Multiple SNPs/Structural’ entries correspond to sequences of length greater than 61, the maximum length of a sequence due to a single SNP with k-mer size of 31 and ‘SNPs’ correspond to sequences of maximum length of 61. (b) Numbers of sequences with alignments to hg38, RefSeq mRNAs and Ensembl exons and coding regions.

### S4 Fig

#### Histograms of sequence lengths in YRI-TSI comparison

Figures show sections of histograms of lengths of sequences associated with (a),(c) YRI and (b),(d) TSI in comparison of YRI and TSI samples. Figures (a), (b) show peaks at 61, the maximum length corresponding to a single SNP with k-mer size of 31. Figures (c), (d) show drop off after 98 which is the maximum length corresponding to two close-by SNPs as 31-mers were assembled using a minimum overlap of 24.

### S5 Fig

#### Histograms of sequence lengths in BEB-TSI comparison

Figures show sections of histograms of lengths of sequences associated with (a),(c) BEB and (b),(d) TSI in comparison of BEB and TSI samples. Figures (a), (b) show peaks at 61, the maximum length corresponding to a single SNP with k-mer size of 31. Figures (c), (d) show drop off after 98 which is the maximum length corresponding to two close-by SNPs as 31-mers were assembled using a minimum overlap of 24.

### S1 Table

#### Variants in *Titin* of differential prevalence in BEB-TSI comparison

Variants in *Titin*, a gene linked to cardiovascular diseases, that were found to be significantly more common in BEB samples compared to TSI samples. The (%) values denote fraction of individuals in the sample with the allele present. The p-values and % values are averaged over *k*-mers constituting the associated sequences.

## Acknowledgments

We thank Faraz Tavakoli, Harold Pimentel, Brielin Brown and Nicolas Bray for helpful conversations in the development of the method for association mapping from sequencing reads using *k*-mers. AR, IH, MBE and LP were funded in part by NIH R21 HG006583. AR was funded in part by Fulbright Science & Technology Fellowship 15093630.

